# Histone deacetylase HDA19 affects cortical cell fate by interacting with SCARECROW in the Arabidopsis root

**DOI:** 10.1101/313791

**Authors:** WQ Chen, C Drapek, DX Li, ZH Xu, PN Benfey, SN Bai

## Abstract

The Arabidopsis root epidermis is a simple model for investigating cell fate specification and pattern formation. In addition to regulatory networks consisting of transcription factors, histone deacetylases are also involved in the cellular patterning process. Here we report HDA19 affects the root epidermal cellular pattern through regulation of cortical cell fate by interacting with SCARECROW. This work reveals two new components in cortical cell specification and uncovers a new facet of SCR function.

## Introduction

The root of Arabidopsis is a simple model that is particularly suitable to the study of cell fate specification and pattern formation. Root tissues are organized as concentric cylinders with roughly three parts: the ground tissue, including an outer layer of cortex and an inner layer of endodermis, encloses the stele, and is surrounded by the outer epidermis ^1^. These tissues are derived from specific groups of initial cells, which, together with the quiescent center (QC), form the stem cell niche at the root tip ^1, 2^.

The root epidermis is composed of two cell types: those that form root hairs (hair cells or H cells) and those that don’t (non-hair cells or N cells). The hair cells normally localize over the cleft between two underlying cortical cells while the non-hair cells lie over a single cortical cell ^3, 4^. Prior to root hair outgrowth, the fate of the two cell types is determined according to their position, and can be distinguished by their different cellular characteristics, such as cytoplasmic density and gene expression ^3, 5, 6^. Previously, it has been reported that a set of transcription factors, including Myb domain proteins WEREWOLF (WER), CAPRICE (CPC), TRIPTYCHON (TRY), AND ENHANCER OF TRIPTYCHON AND CARPRICE1 (ETC1); bHLH domain proteins GLABRA3 (GL3) and ENHANCER OF GLABRA3 (EGL3); WD-repeat protein TRASNPARENT TESTA GLABARA (TTG); the homeo-domain protein GLABRA2 (GL2), and others, are responsible for the cellular patterning of the epidermis through a complicated interaction network ^7–12^. Two membrane-localized receptor-like kinases, SCRAMBLED (SCM) and BRASSINOSTEROID INSENSTITIVE 1 (BRI1), are involved in the cellular patterning by mediating an unknown positional signal derived from the underlying cortical cells ^13, 14^. In addition to transcription factors, histone modifications, especially histone acetylation were found to enhance the robustness of the transcription factor network ^15–17^ and regulate kinase gene expression ^18^.

Except for the role of JACKDAW (JKD), little is known about how cortical cell differentiation plays a role in the cellular patterning of the root epidermis ^19^. A single cortex layer is initiated from the cortex/endodermis initial (CEI) cells. Both cortex and endodermis are derived after two successive asymmetric cell divisions of the CEI. The GRAS family transcription factors SCARCROW (SCR) and SHORT-ROOT (SHR) play critical roles in ground tissue patterning and identity maintenance ^20, 21^. SCR is predominantly expressed in the QC, the CEI and the endodermis. SHR moves from the stele into the endodermis where it is sequestered in the nucleus by SCR ^22–24^. The interaction between SCR and SHR determines CEI identity, promotes its asymmetric cell division and specifies the fates of cortex and endodermis by forming a cell-type-specific regulatory network with other transcription factors such as JKD, MAGPIE (MGP), NUTCRACKER (NUC) and other BIRD factors ^25–27^. As the root ages, the endodermis undergoes another asymmetric cell division to give rise to a second layer of cortex, termed middle cortex (MC) ^28, 29^. Interestingly, SCR suppresses this second cell division; in contrast to its promoting effect for the CEI division ^30^. It has been suggested that the two opposite functions of SCR are achieved by interacting with different partners through different domains ^31^.

Previously, it was demonstrated that an inhibitor of histone acetylation, trichostain A (TSA) can induce hair cell differentiation at non-hair positions ^15^ . We carried out a systematic screen to clarify which members of the histone acyltransferase (HAT) and histone deacetylase (HDAC) family are involved in cellular patterning of the root epidermis ^32^. In addition to HDA18 and HDA6, mutations in *HDA19* also exhibit an altered cellular pattern of the root epidermis ^16, 18, 32^. Here we report that HDA19 functions primarily in differentiation of the cortical cells through interaction with SCR in the CEI cells. The altered cellular patterning of root epidermis observed in the *hda19* mutant is thus derived from altered differentiation of the cortex. These findings demonstrate that the HDA19 enhances the function of SCR in its role in regulating ground tissue differentiation in the Arabidopsis root.

## Results

### HDA19 is required for cellular patterning of root epidermis and cortex

In a screen for altered cellular patterning of the root epidermis caused by mutations of HDAC family members, we found an altered cellular pattern of the epidermis in the *hda19* mutant ^32^. In addition to the significant increase of both ectopic H cells at the N positions and ectopic N cells at the H positions in the epidermis (Table 1), cortex cell number was greatly increased at 8 days after sowing, and the middle cortex layer had already appeared in most *hda19* roots (95% relative to 0% in Col) (Fig. 1a, b). There was also a slight increase in endodermis cell number (Table. 1). We complemented the mutation with *HDA19pro:HDA19-EGFP* in *hda19* (Fig. 1d, Table. 1) and found GFP signal in all the cell layers of the root tip (Fig. S1d).

**Fig1.**
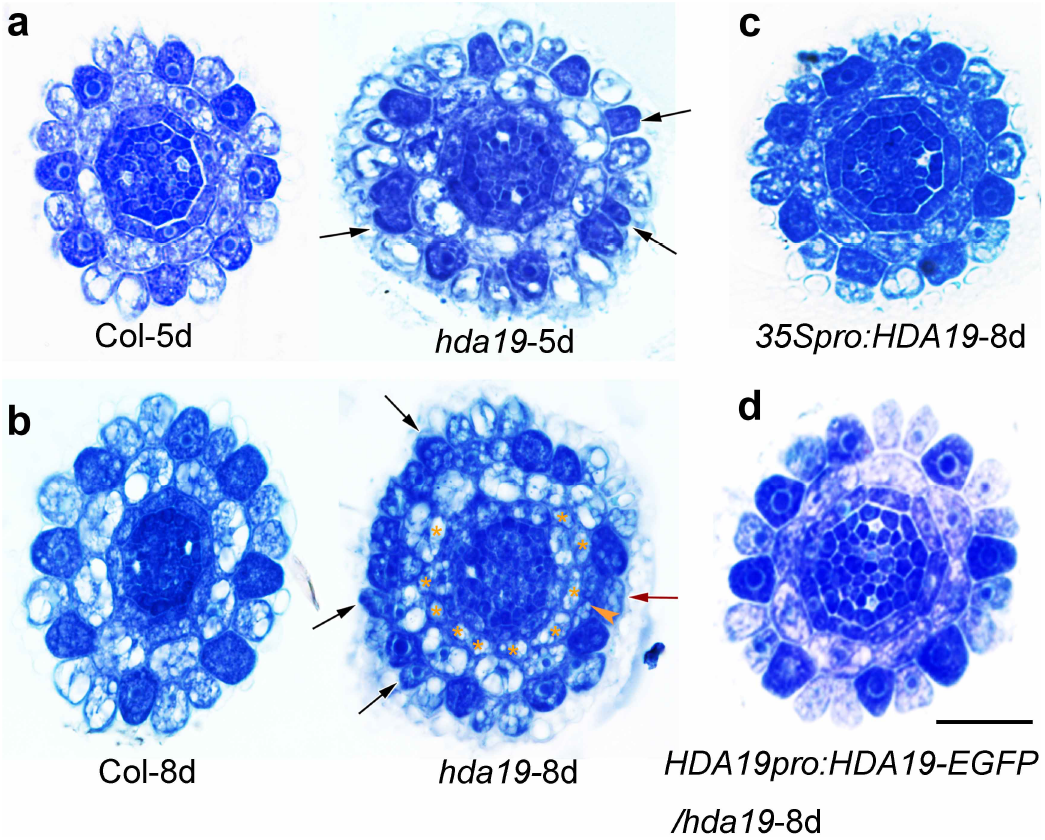
The *hda19* mutant has defects in root epidermal cell and ground tissue patterning. Cross section images of toluidine blue stained root tips of (a) wild type (Col) and *hda19* mutant at 5 day-old and (b) 8 day-old; (c) *35Spro:HDA19* (over-expression line) at 8 day-old and (d) *HDA19pro:HDA19-EGFP/hda19* (complementation line) at 8 day-old. Black arrows indicate darkly stained H cells at N positions and red arrows indicate lightly stained N cells at H positions. Orange asterisks indicate the additional layer of the ground tissue and orange arrowhead indicates the additional anticlinal cell division of cortical cell. Scale bar = 20 μm.

**Table 1.** Quantification of ectopic epidermal cell differentiation and abnormal ground tissue cell number in the root tips of 8 day-old seedlings of wild type (Col), *hda19* mutant, *HDA19* over-expressing line and HDA19-EGFP driven by *HDA19* promoter and tissue-specific promoters.

| Sample | Ratio of root hair (%) | N cell at H position (%) | H cell at N position (%) | Cortex cell number | Endodermis cell number | Ratio of MC (%) | n |
| --- | --- | --- | --- | --- | --- | --- | --- |
| Col | 42.3±2.2 | 0.8±1.3 | 2.0±2.8 | 8.0±0.0 | 8.0±0.1 | 0 | 17 |
| *hda19* 5 day-old | 47.4±3.1 <sup>a</sup> | 8.1±6.1 <sup>a</sup> | 17.2±4.9 <sup>a</sup> | 9.4±1.4 <sup>a</sup> | 8.3±0.4 | 46.2 | 12 |
| *hda19* 8 day-old | 46.7±5.4 | 21.9±8.5 <sup>a</sup> | 17.9±7.9 <sup>a</sup> | 13.6±1.4 <sup>a</sup> | 9.4±0.8 <sup>a</sup> | 95 | 20 |
| *35Spro:HDA19* | 42.3±2.4 | 0±0 | 1.8±2.7 | 8.0±0.1 | 8.0±0.0 | 0 | 23 |
| *HDA19pro:HDA19-EGFP/hda19* | 43.3±3.7 | 2.7±4.2 <sup>b</sup> | 1.9±2.9 <sup>b</sup> | 8.5±0.8 <sup>b</sup> | 8.5±0.6 | 11.5 | 26 |
| *WERpro:HDA19-EGFP/hda19* | 40.6±3.7 | 17.8±9.2 <sup>a</sup> | 8.0±4.0 <sup>a b</sup> | 10.4±1.7 <sup>a</sup> | 8.8±0.6 <sup>a</sup> | 50 | 20 |
| *JKDpro:HDA19-EGFP/hda19* | 43.3±3.1 | 4.5±4.3 <sup>b</sup> | 3.9±3.9 <sup>b</sup> | 8.4±0.5 <sup>b</sup> | 8.3±0.5 | 19 | 26 |
| *SCRpro:HDA19-EGFP/hda19* | 43.9±4.0 | 1.5±2.0 <sup>b</sup> | 7.2±5.2 <sup>a b</sup> | 8.5±0.7 <sup>b</sup> | 9.0±0.8 <sup>a</sup> | 12.5 | 32 |
| *SHRpro:HDA19-EGFP/hda19* | 47.7±3.6 <sup>a</sup> | 14.6±4.8 <sup>a</sup> | 15.9±4.8 <sup>a</sup> | 9.3±1.2 <sup>a b</sup> | 8.3±0.5 | 67 | 21 |

Values indicate mean±SD (a: differs significantly from the wild type Col; b: differs significantly from *hda19* mutant, p<0.01. Student's *t* test). 8 day-old seedlings are used except *hda19* 5 day-old. n means the number of seedlings examined.

To determine if HDA19 directly affects the cellular patterning of the root epidermis, we examined the expression of known patterning genes using quantitative reverse transcription PCR (RT-qPCR). In *hda19*, *CPC*, *TRY*, and *SCM* were down-regulated, while *ETC1* and *MYB23* were up-regulated (Fig. 2a). Using marker lines, we found no changes in the *WERpro:GFP* expression pattern (Fig. 2b). In lines expressing *GL2pro:GFP* and *CPCpro*:*GUS*, several cells at the N position had undetectable *GL2pro:GFP* signal and ectopically expressed *CPCpro*:*GUS* (Fig. 2c, d), indicating that these cells adopted the H cell fate. Interestingly, the expression from *SCMpro*:*GUS* was dramatically decreased in the root tip, while the signal in the hypocotyl was unchanged as compared to wild type (Fig. 2e). Down-regulation of *SCM* expression in the root was confirmed by RT-qPCR (Fig. 2a). Since SCM is a membrane receptor-like kinase important for sensing an unknown positional signal derived from the cortex, the effect of HDA19 on the regulation of *SCM* expression implies that HDA19 may act upstream of this regulatory network.

**Fig2.**
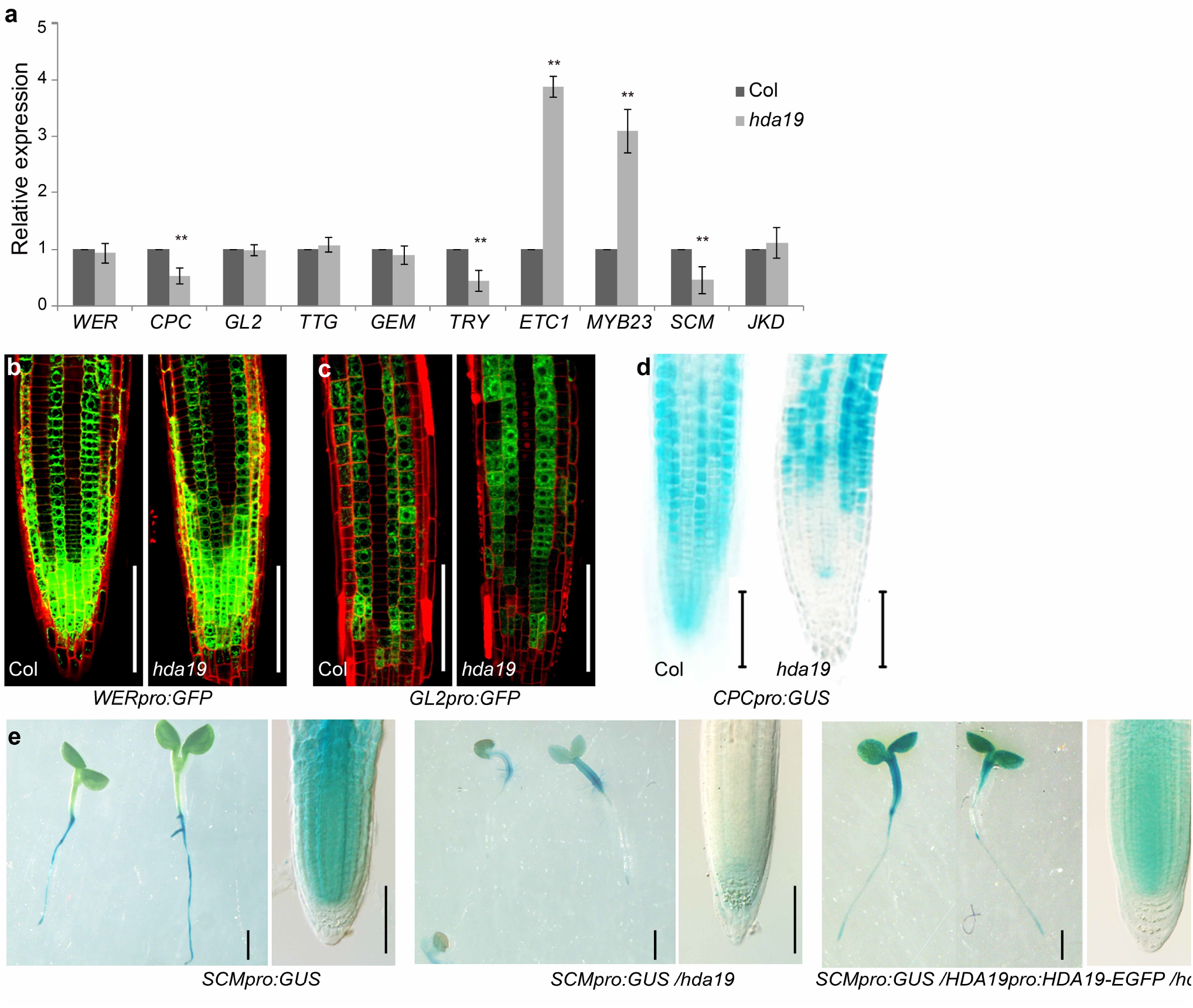
*HDA19* affects the expression of root epidermal patterning genes. (a) Expression level of genes in 8 day-old root tips of *hda19* determined via qRT-PCR. The values over wild type (Col) are shown as means ± standard deviation (SD, *P < 0.05, **P < 0.01; Student’s *t* test). (b - c) Confocal microscopy images of the root tip (epidermal view) of (b) *WERpro:GFP* and (c) *GL2pro:GFP* expression pattern in wild type and *hda19* background, respectively. Scale bar = 100 μm. (d) *CPCpro:GUS* expression in 5 day-old root tip in wild type and *hda19* background, respectively. Scale bar = 100 μm. (e) *SCMpro:GUS* expression in 5 day-old seedlings (left) and root tip (right) in wild type, *hda19* and *HDA19pro:HDA19-EGFP /hda19* background, respectively. Scale bar left = 1mm, scale bar right = 100 μm.

To determine if HDA19 can bind directly to the patterning genes whose expression is altered in *hda19*, we performed chromatin immunoprecipitation and found that none of the tested regions of the patterning genes was enriched (Fig. S2a, b). However, global elevation of histone H3 and H4 acetylation levels on these genes was observed (Fig. S2c, d), both when these genes were up and down-regulated in *hda19*. To rule out an effect of HDA19 on the proteins encoded by the patterning genes, we carried out a yeast two-hybrid assay. No relevant protein-protein interactions were found in this assay (Fig. S3). These results suggest that HDA19 regulates expression of these genes in an indirect manner.

### HDA19 affects cortical cell fate required for epidermal differentiation

As our data indicate that the altered cellular pattern of the root epidermis in *hda19* does not result from direct regulation by HDA19, we hypothesized that HDA19 acts through control of cortex differentiation. To determine the origin of the additional cortex cell layer we examined the initial cells and found an ectopic periclinal cell division (Fig. 1a; Fig. 3a-d, indicated by asterisks), which explained the significant increase of total cell number of cortex and endodermis in *hda19* (Table. 1). Between 5 and 8 days, there were also additional anticlinal cortical cell divisions in *hda19*, and the percentage of N cells at H positions rose from an average of 8.1% to 21.9%. A likely scenario is that cells at the newly emerged H position between two cells produced by the ectopic anticlinal cortical cell division failed to change fate accordingly (Fig. 1b, indicated by arrowhead and red arrow). To test this hypothesis, we needed to confirm that the cells produced by the additional periclinal divisions are cortical cells. We introduced the cortex and endodermis specific marker lines *CO2pro:NLS-YFP* and *En7pro:NLS-YFP* into *hda19*. Strikingly, the signal of the cortex-specific marker *CO2pro:NLS-YFP* in *hda19* was greatly decreased, with the signal observed only randomly in a few cortical cells (Fig. 3a). This result was confirmed by RT-qPCR (Fig. 3e). By contrast, the endodermis specific marker *En7pro:NLS-YFP* was normally expressed in *hda19* (Fig. 3b, e). We also examined the expression pattern of *SCRpro:GFP-SCR* and *SHRpro:SHR-GFP* in *hda19* and found no obvious changes (Fig. 3c, d). These results indicate that HDA19 mainly affects the identity of the root cortex, but not the endodermis.

**Fig3.**
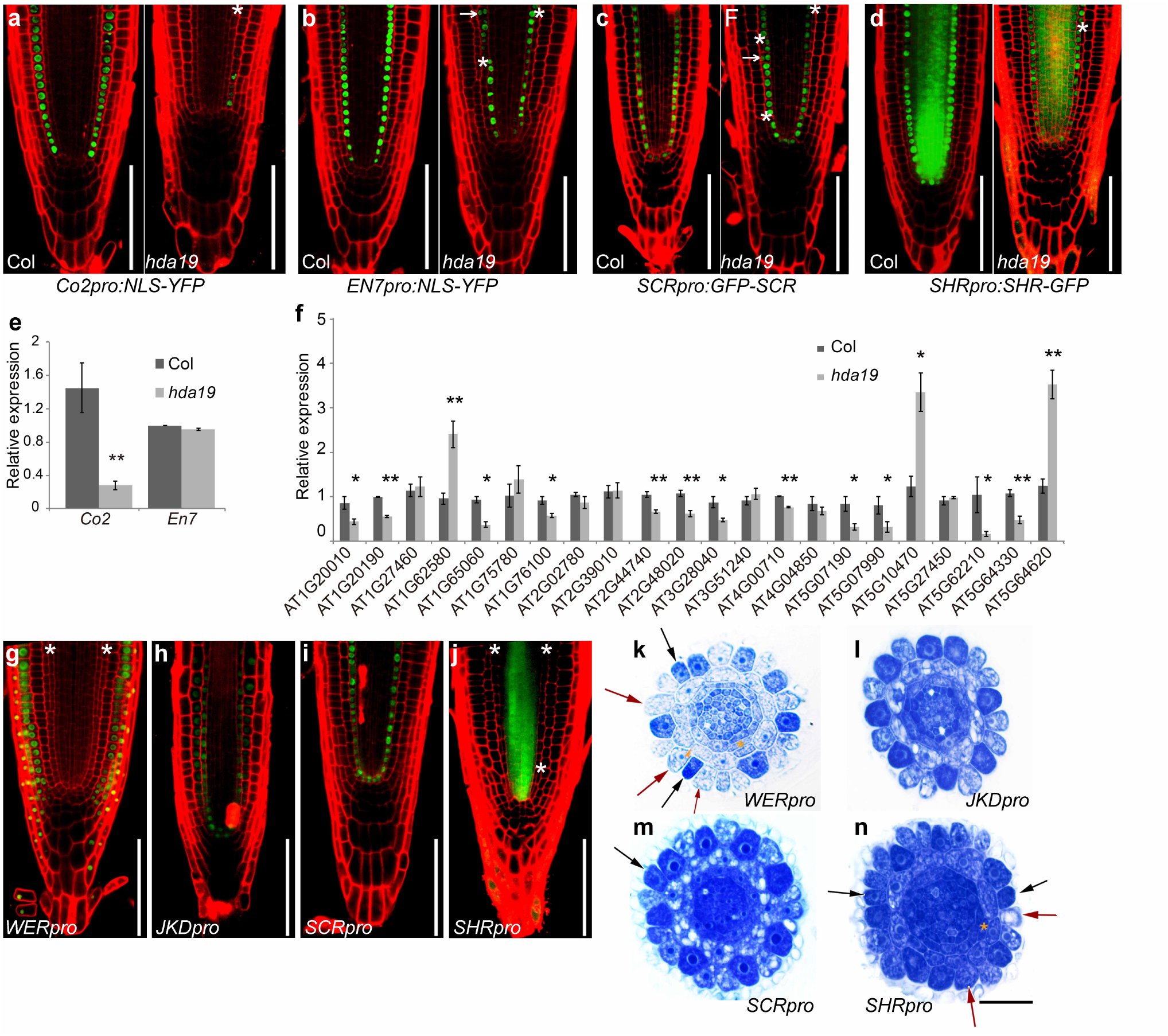
*HDA19* affects root cortex gene expression and regulates proper epidermal differentiation though its action in the ground tissue. (a - d) Confocal microscopy images of YFP or GFP signal in root tips of 8 day-old seedlings. White asterisks indicate the additional middle layer of the ground tissue. White arrows indicate the YFP or GFP signal remaining in the out layer of daughter cells after cell division. Scale bar = 100 μm. (e) Expression level of *Co2* and *En7* genes in 8 day-old root tips of *hda19* determined via qRT-PCR. (f) Expression level of selected cortex-specific expression genes in 5 day-old root tips of *hda19* determined via qRT-PCR. Error bar represents SD value from three biological replicates (*P < 0.05, **P < 0.01; Student’s *t* test). (g - j) Confocal microscopy images of HDA19-EGFP 8 day-old root tip expression driven by tissue-specific promoters in *hda19*. Asterisks indicate the additional layer of ground tissue. Scale bar = 100 μm. (k - n) Cross section and toluidine blue stained root tips of 8 day-old seedlings of HDA19-EGFP driven by tissue-specific promoters in *hda19*. Black arrows indicate the darkly stained H cells at N positions and red arrows indicate the lightly stained N cells at H positions. Orange asterisks indicate the additional layer of ground tissue. Scale bar = 20 μm.

To confirm the findings based on marker line analysis, we examined gene expression levels by RT-qPCR. Among 82 genes specifically expressed in meristematic cortex (^33^; http://bar.utoronto.ca/efp/cgi-bin/efpWeb.cgi), 4 GO categories were identified, including membrane, glycoprotein, nucleotide-binding and signaling genes. We selected 22 meristematic cortex genes from these 4 groups (Table. S1) and found that 12 genes were down regulated and 3 upregulated in *hda19* (Fig. 3f). These results indicate that HDA19 affects cortical cell identity.

### HDA19 acts in the ground tissue to regulate epidermal cell fate

To test if HDA19 controls epidermal cell fate by acting in the cortex, we drove *HDA19* with the tissue specific promoters *WERpro* (epidermis-specific), *JKDpro* (ground tissue-specific), *SCRpro* (endodermis-specific) and *SHRpro* (vascular tissue-specific) in *hda19*. As mentioned above, *HDA19pro:HDA19-EGFP/hda19* fully complemented the mutant phenotype (Fig. 1d), demonstrating the functionality of the *HDA19-EGFP* construct. We found that HDA19-EGFP in the epidermis under the *WERpro* failed to restore either epidermis or ground tissue patterning (Fig. 3 g, k). HDA19-EGFP expression in the ground tissue under *JKDpro* was able to fully restore both epidermal differentiation and the ground tissue cell division pattern (Fig. 3h, l). Unexpectedly, endodermis-specific expression of HDA19-EGFP under *SCRpro* could also restore the cortical cell division pattern and partially rescue the epidermal phenotype of *hda19* (Fig. 3i, m). However, expression of HDA19-EGFP in the vascular tissue under *SHRpro* still exhibited abnormal epidermal and ground tissue patterning (Fig. 3 j, n). We also examined the expression pattern of *SCRpro:GFP-SCR* and *SHRpro:SHR-GFP* in *hda19* and found no obvious changes (Fig. 3c, d), indicating that the CEI and endodermis identity are intact in *hda19*. Taken together, our results demonstrate that HDA19 affects cortex cell fate and gene expression. Moreover, HDA19 expressed in the ground tissue is able to non-cell-autonomously regulate epidermal patterning.

### HDA19 interacts with SCR in the CEI

To determine how HDA19 affects cortex differentiation we asked if HDA19 can interact with known factors involved in ground tissue differentiation. Using a yeast-two-hybrid assay (Y2H), we found that HDA19 can interact with SCR and MGP, but not with SHR or JKD (Fig. 4a; Fig. S4). Also, HDA19 could not indirectly interact with SHR or JKD using SCR or MGP as a bridge in a yeast-three-hybrid assay (Fig. S5). To identify the interacting domains, we performed a Y2H with truncated fragments of HDA19 and SCR. Interaction took place between the HDAC domain of HDA19 and the ND domain (N-terminal variable domain, known to suppress ectopic asymmetric cell division) of SCR ^30, 34^ (Fig. 4b, c; Fig. S4a, b). We confirmed the interaction of HDA19 and SCR *in planta*, using BiFC (bimolecular florescence complementation) and transient expression (Fig. 4d).

**Fig4.**
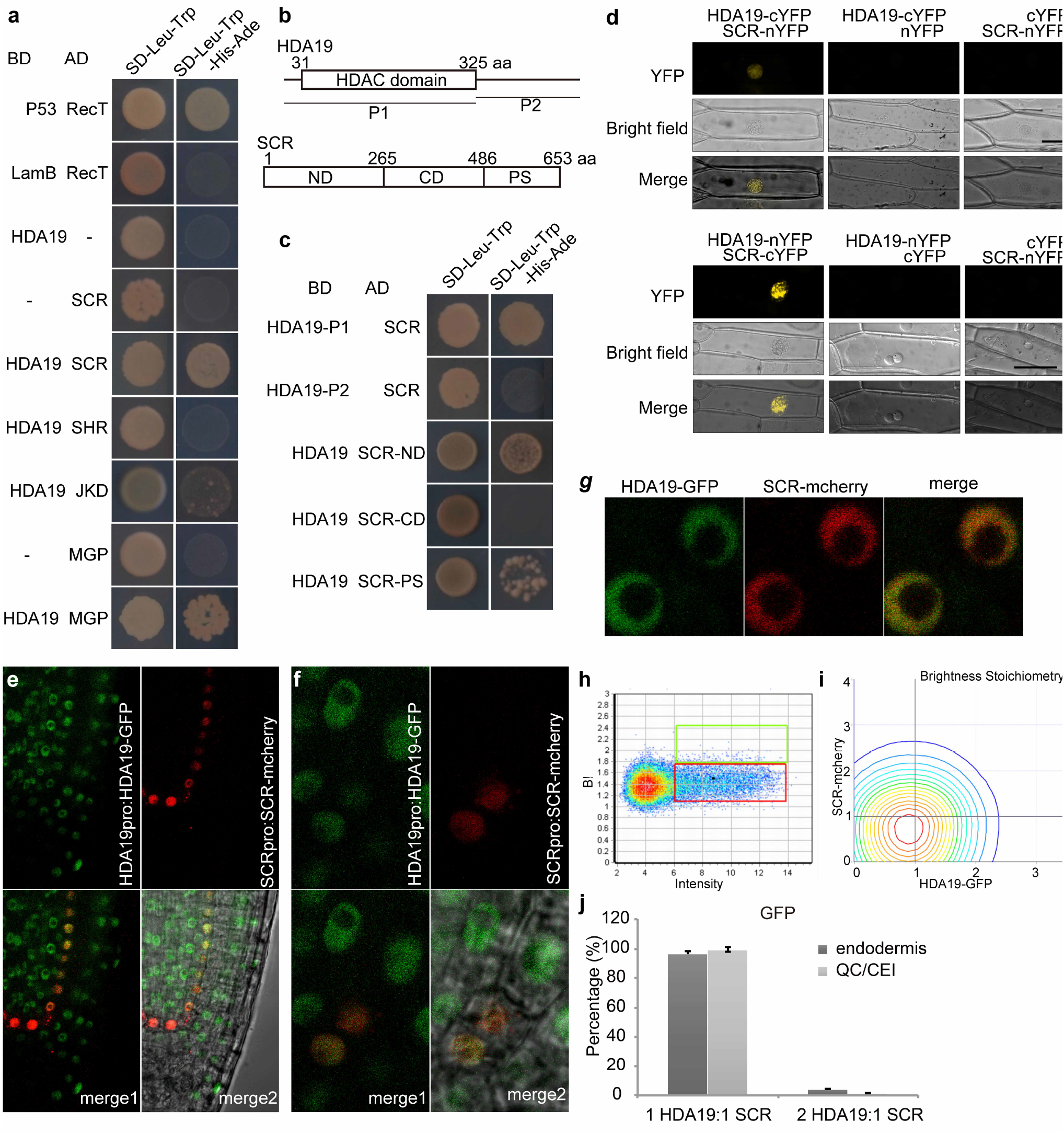
HDA19 interacts with SCR. (a) Yeast-two-hybrid analysis of protein interactions between HDA19 and some proteins important to ground tissue development. (b) Molecular structure of HDA19 protein and SCR protein, and their fragments used in the yeast-two-hybrid assay. (c) Yeast-two-hybrid analysis of the interaction between fragments of HDA19 and SCR protein, and between HDA19 protein and fragments of SCR, respectively. (d) BiFC assay of HDA19 and SCR protein interaction in onion epidermis as the living cell. nYFP, N terminus of YFP; cYFP, C terminus of YFP. Scale bar = 50μm. (e) Confocal images of longitudinal root sections in *HDA19pro:HDA19-GFP* and *SCRpro:SCR-mCherry* line. (f) Confocal images of CEI/CEID cells in *HDA19pro:HDA19-GFP* and *SCRpro:SCR-mCherry* line. (g) Confocal images of HDA19-GFP/SCR-mCherry nuclei used for cross N&B analysis. (h) Brightness vs Intensity plot for HDA19pro:HDA19-GFP in the meristem. The red and green boxes represent the HDA19-GFP monomer and homodimer, respectively. (i) Stoichiometry histogram from cross N&B analysis of *HDA19pro:HDA19-GFP* and *SCRpro:SCR-mCherry*. (j) Average percentages of 1:1 and 2:1 HDA19-GFP and SCR-mCherry interaction in endodermis and QC/CEI cells. Error bar represents SD (n=5).

To determine how SCR, an endodermis-specifically expressed protein interacts with HDA19 we carried out Fluorescence Correlation Spectroscopy (FCS) to see where the two proteins interact *in vivo*. FCS is a non-invasive technique that measures fluctuations of fluorescently-tagged proteins *in vivo* to determine dynamics of protein binding and movement. Specifically, we used an FCS technique called cross-correlation Number and Brightness to determine the binding and stoichiometric ratio of the SCR-HDA19 complex ^24, 35, 36^. To do this, we generated a *HDA19pro:HDA19-EGFP x SCRpro:SCR-mCherry* (Fig. 4e-g) and measured cross-correlation of EGFP and mCherry brightness in the QC, CEI and endodermis (Fig. 4h, i). We found SCR and HDA19 interact in vivo and bind primarily (98.8%) in a 1:1 to ratio (Fig. 4j).

### HDA19 binds to *SCR* promoter and SCR target genes and affects their expression

To determine if HDA19 functions through interaction with SCR, we performed ChIP-PCR using a GFP antibody in the *HDA19pro:HDA19-EGFP/hda19* line. We found enrichment of the promoter sequence of SCR as well as the promoter regions of the SCR target genes, *MGP*, *NUC*, *RLK* and *BR6OX2* (Fig. 5a-g). To determine if the binding of HDA19 to the SCR target genes is dependent on SCR, we examined binding of *HDA19pro:HDA19-EGFP/hda19* in a *scr-3* background and found increased enrichment of several regions upstream of *SCR*, *MGP* and *BR6OX2* (Fig. 5b-g). This suggests that not only can HDA19 directly bind to the chromatin upstream of *SCR* and its target genes, but also SCR interferes with the binding of HDA19 protein to some of its target genes.

**Fig5.**
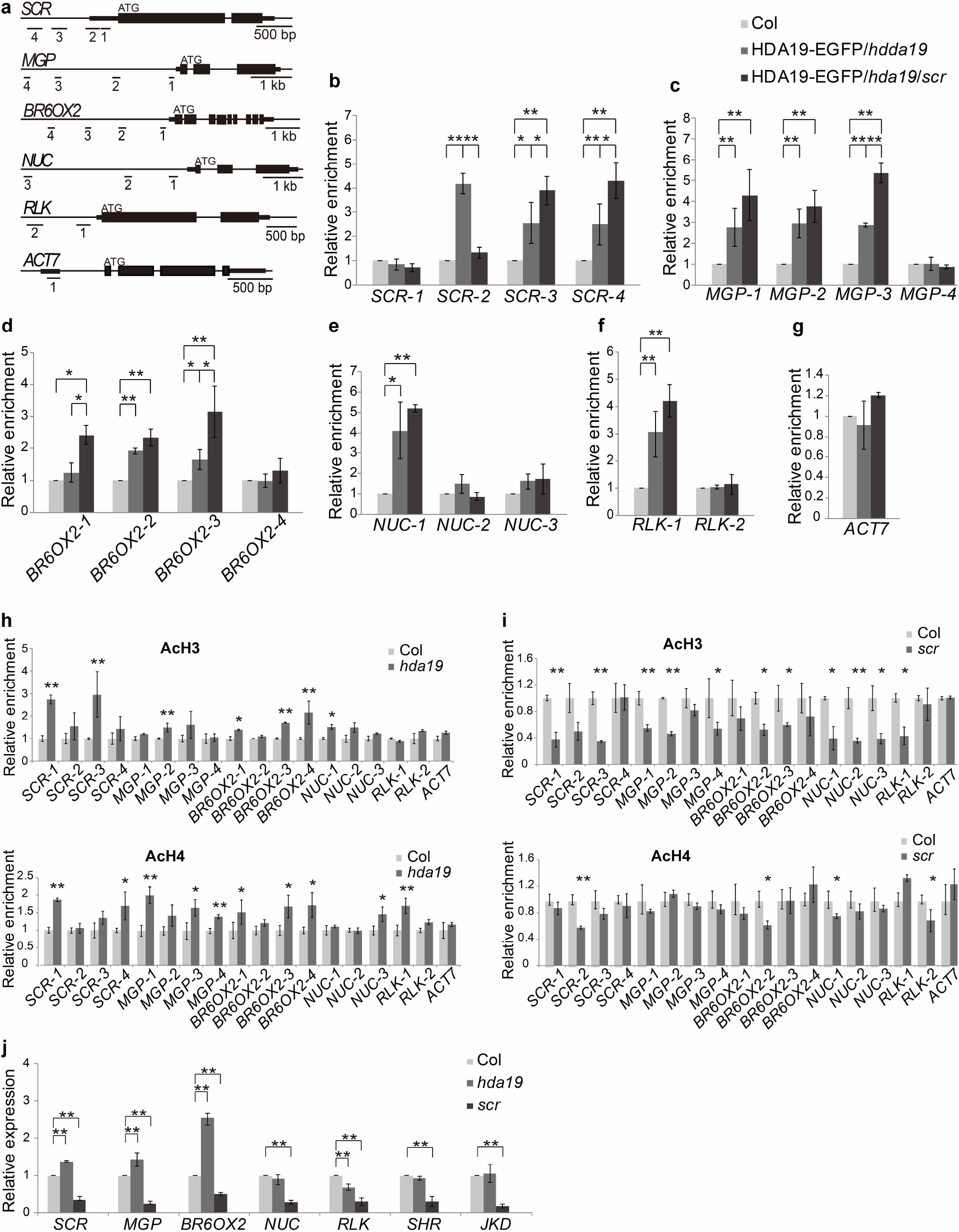
HDA19 binds to SCR’s target genes and regulates their expression. (a) Schematic of the *SCR* gene and SCR’s target genes. Regions selected for ChIP analysis are underlined. (b - g) ChIP-qPCR analysis of HDA19 binding via GFP antibody in *HDA19pro:HDA19-EGFP/hda19* (HDA19-GFP*/hda19)* and *HDA19pro:HDA19-EGFP/hda19/scr* (HDA19-GFP*/hda19/scr*). The values over wild type (Col) are shown as means±SD (*P < 0.05, **P < 0.01; Student’s *t* test). (h - i) ChIP-qPCR analysis of histone H3 and H4 acetylation level changes in *hda19* (h) and *scr-3* (i) compared to wild type (Col). Error bar represents SD value (*P < 0.05, **P < 0.01; Student’s *t* test). (j) Expression of target genes in the 5 day-old root tip of *hda19* and *scr-3* compared to Col determined via qRT-PCR. The values over Col are shown as means±SD (*P < 0.05, **P < 0.01; Student’s *t* test).

To test if HDA19 functions through its histone deacetylase activity, which was confirmed *in vitro* as reported in ^37, 38^, we examined acetylation levels of H3 and H4 at the promoter regions of the HDA19 bound genes. We found that acetylation levels of common HDA19 and SCR bound genes were increased in *hda19* (Fig. 5h), indicating regulation by deacetylation activity of HDA19. In contrast, the acetylation levels of H3 and H4, especially H3 were significantly decreased for the HDA19 and SCR bound genes in *scr* (Fig. 5i). This is consistent with the finding that SCR may interfere with binding of HDA19 to its target genes.

We next determined the expression levels of the target genes in *hda19* and *scr*. Consistent with previous reports ^22^, expression of *MGP*, *NUC*, *RLK* and *BR6OX2* were greatly down regulated in *scr*. In *hda19*, expression of *SCR*, *MGP* and *BR6OX2* were up regulated (Fig. 5j), suggesting that HDA19 plays a role in modifying the transcriptional activity of SCR for some of its target genes. As expected, no such effect was observed in the expression of *SHR* or *JKD* (Fig. 5j).

Taken together, we conclude that HDA19 interacts with SCR, probably in the CEI and endodermis, and affects the expression of *SCR* and some SCR target genes.

### SCR and its target genes are involved in Arabidopsis root epidermal differentiation

The finding that SCR interacts with HDA19 at the ND domain led to the hypothesis that the ND domain is important for cortex differentiation. Detailed examination of the cortical cell division in *SCRΔND* lines revealed ectopic anticlinal cortical cell divisions resulting in increased cortical cell numbers (Fig. 6c; Table. 2). In addition, expression of *MGP* and *BR6OX2* was significantly up regulated in *SCRΔND* roots (Fig. S6). *SCRΔND* also had an altered epidermal differentiation pattern, with both ectopic H cells at N position and N cells at H position (Fig. 6c; Table. 2). *SCM* expression in *SCRΔND* root tips was slightly down regulated while it was highly up regulated in *scr-3* (Fig. S7f). The phenotype and gene expression results in the *SCRΔND* lines were similar to those in *hda19* indicating that *SCR* can regulate epidermal cell specification and cortex cell division.

**Fig6.**
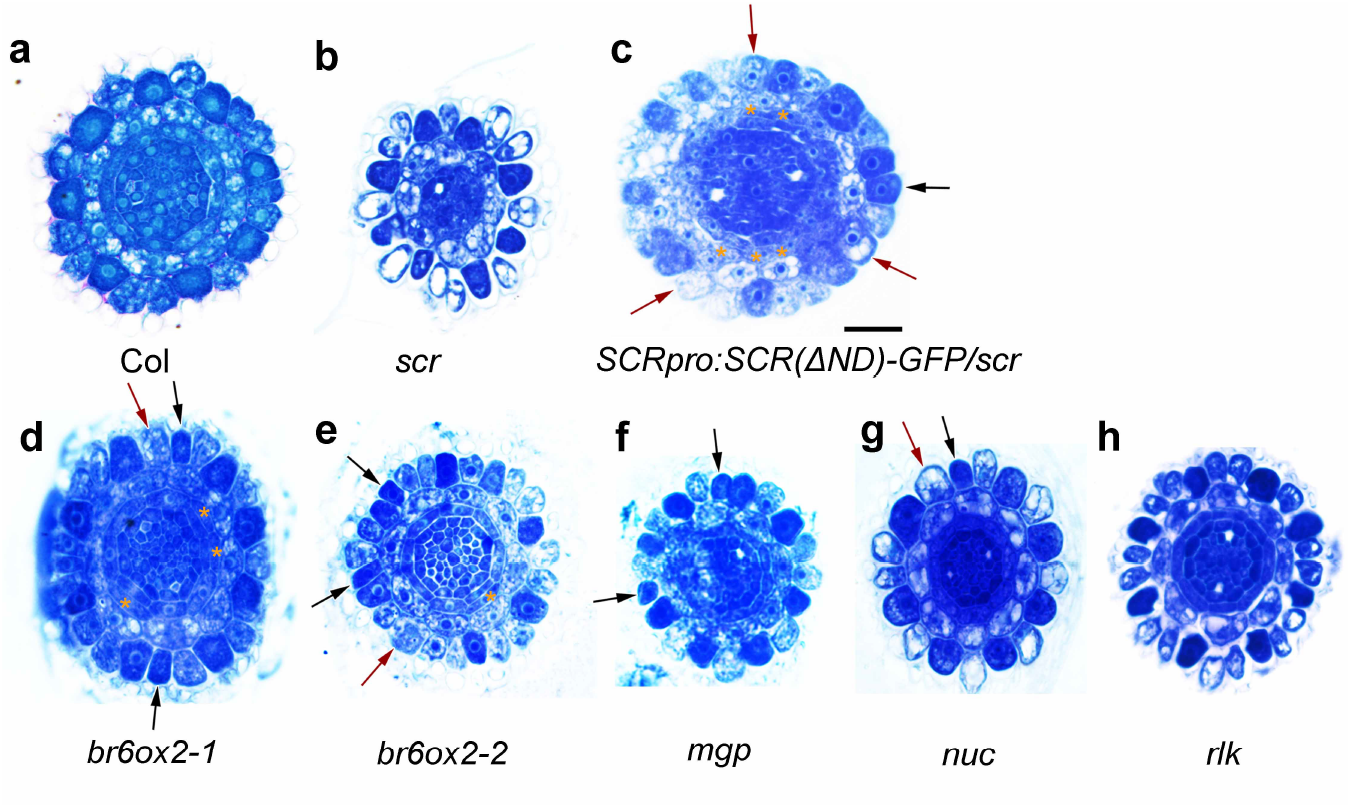
Mutant alleles of *scr*, *SCRΔND* and its target genes have defects in root epidermal cell and ground tissue patterning. Cross section and toluidine blue stained root tips of 8 day-old seedlings in (a) wild type, (b) *scr-3*, (c) SCRpro:SCR(∆ND)-GFP/*scr-4*, (d) *br6ox2-1*, (e) *br6ox2-2*, (f) *mgp*, (g) *nuc*, (h) *rlk*.. Black arrows indicate the darkly stained H cells at N position. Red arrows indicate the lightly stained N cells at H position. Orange asterisks indicate the additional layer of ground tissue. Scale bar = 20 μm.

**Table 2.**
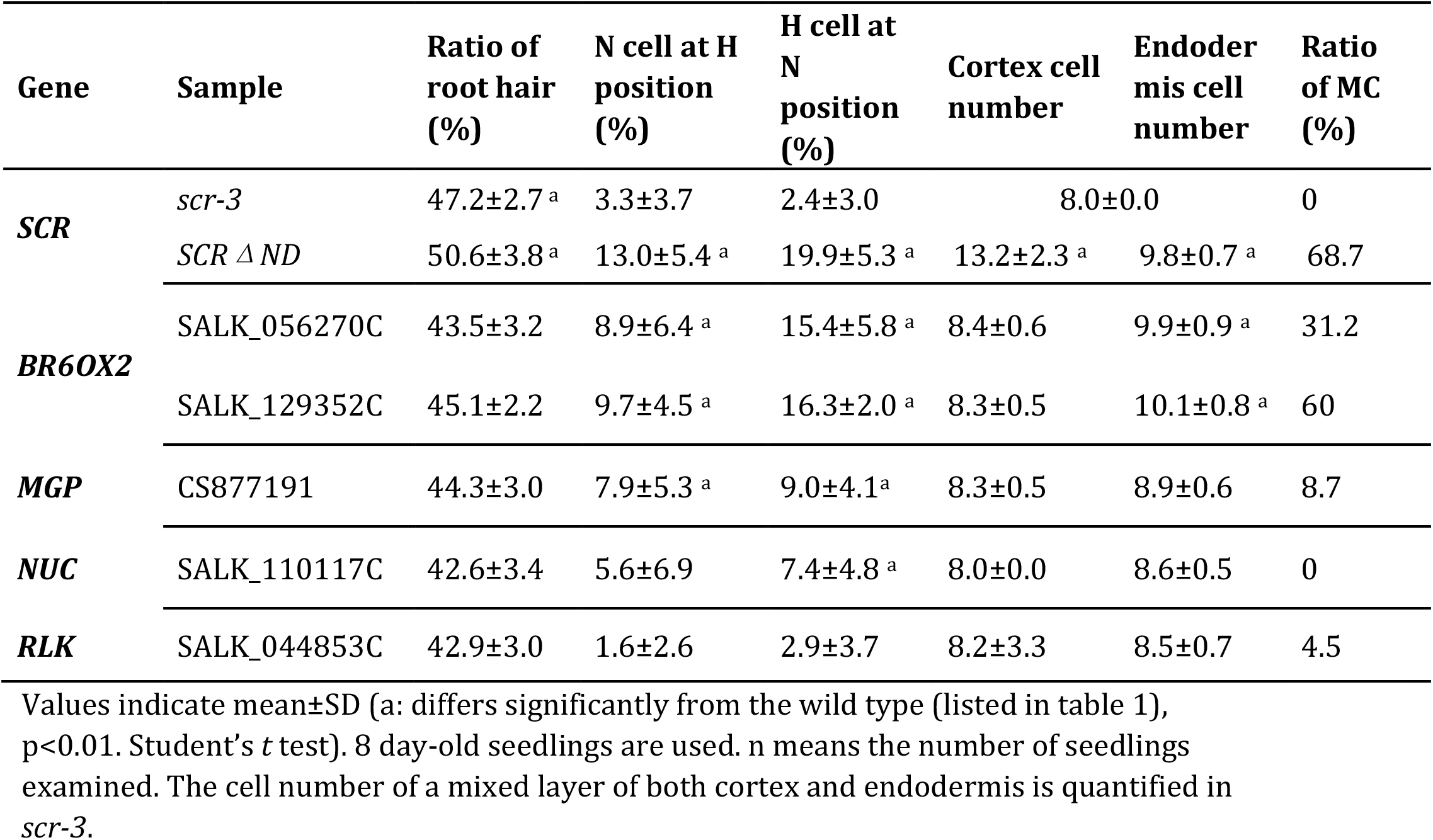
Quantification of ectopic epidermal cell differentiation and abnormal ground tissue cell number in the root tips of 8 day-old seedlings of *scr-3*, *SCRΔND* and mutant alleles of SCR target genes.

We also found two T-DNA insertion mutant lines of *BR6OX2* that exhibited disturbed epidermal patterning and ectopic periclinal cell division in the ground tissue, similar to *hda19* and *SCRΔND* (Fig. 6d, e; Table. 2). In addition, *SCM* expression was significantly reduced in *br6ox2* and *mgp* (Fig. S7c-f). *BR6OX2* encodes an enzyme that catalyzes the last reaction of brassinolide (BL) synthesis suggesting that BL plays a role in the positional signal from ground tissue to control epidermal differentiation.

### Discussion

We have found that the altered cellular pattern of the root epidermis in the *hda19* mutant is derived from abnormal cortex differentiation. HDA19 achieves this through interaction with SCR and by directly binding SCR target genes. Interaction between HDA19 and SCR is mainly localized to the CEI, suggesting the possibility of preprograming of cortex cell fate in its progenitor cells. This work not only provides a molecular mechanism underlying the phenotype of *hda19*, but also reveals a new function of SCR.

Arabidopsis root tissues are organized roughly into three parts: epidermis, stele, and ground tissue in between. The ground tissue is initiated from the CEI at the root tip and further differentiated into an outer layer of cortex and inner layer of endodermis. While multiple genes are known to be involved in the regulation of epidermal cellular pattering ^39–41^, little is known about the control of cortex identity except for the role of *JACKDAW* (*JKD*) ^19^. The present work adds two additional players, HDA19 and SCR, to the network regulating cortex differentiation. The effect of HDA19 on cortex differentiation through interaction with SCR in the CEI suggests a novel regulatory pathway, which may be parallel to that of JKD.

HDA19 is a global regulator of transcription in response to a variety of development processes and stress, including seed germination, root apical defects, leaf shape and flower structure ^42–45^. In contrast, SCR is relatively specific in its role in endodermal cell fate specification. Thus, HDA19 may acquire its spatiotemporal specificity in its function in cortex cell fate determination through interaction with SCR. It is interesting that a point mutation in SCR that causes a premature stop at codon 460 and loss of the C-terminal PS domain, did not exhibit abnormal cortex or epidermis, while deletion of its ND did ^30, 46^. This can be explained by the interaction of SCR and HDA19 at SCR’s ND. This finding provides new insight into the molecular mechanism of SCR regulation.

Brassinosteroids are required to maintain the cellular pattern of the Arabidopsis root epidermis ^14^. Our finding that mutation of the BR synthesis gene *BR6OX2*, a SCR and HDA19 target, results in an abnormal epidermal and cortex pattern (Fig. 6f) suggests that BR signaling originates in cortical cells and is regulated by HDA19 and SCR.

## Materials and Methods

### Plant Material and Growth Conditions

The *Arabidopsis* (*Arabidopsis* thaliana) SALK and CS series of T-DNA insertion lines of *hda19* (SALK_139445), *br6ox2-1* (SALK_129352C), *br6ox2-2* (SALK_065270C), *mgp* (CS877191), *nuc* (SALK_110117C) and *rlk* (SALK_044853C) mutant alleles were purchased from the Arabidopsis Biological Resource Center (http://www.arabidopsis.org/). The following mutants and reporter lines were previously described: *scr-3* ^20^, *SCRpro:GFP-SCRΔND/scr-4* line ^30^, *Co2pro:NLS-YFP*, *En7pro:NLS-YFP*, *SCRpro:GFP-SCR*, *SHRpro:SHR-GFP* ^22^; *WERpro:GFP* ^9^, *GL2pro:GFP* ^47^, *CPCpro:GUS* ^48^ and *SCMpro:GUS* ^13^. Reporter lines were introduced into *hda19* or transgenic plants by crossing. Homozygous lines were obtained by molecular genotyping, phenotype analysis, and reporter gene expression verification. Seeds were sterilized and placed on Murashige and Skoog (MS) medium solidified with 0.8%-1.0% (m/v) agar. Plants were grown under 22°C chamber with a period of 16 h light /8 h dark.

### Construction of Plasmids and Transgenic Plants

To over express *HDA19*, the coding sequence of *HDA19* (without the stop codon) was cloned after CaMV35S promoter and fused in frame to Flag epitope using the pFPZP110 vector. To construct *HDA19pro:HDA19-EGFP*, a 2562 bp fragment of *HDA19* genomic sequence (from ATG but without the stop codon) was introduced into the pEGAD-EGFP vector fused in frame of the EGFP epitope to get HDA19-EGFP construct. Then, a 1452 bp upstream region of the *HDA19* (including the intergenic sequence, 5’ UTR and the first intron) was first introduced into the pEGAD-EGFP vector to replace the CaMV35S promoter and a 385 bp downstream fragment (including the intergenic sequence and 3’ UTR) was inserted at the end of EGFP epitope. For tissue specific expression of *HDA19* constructs, a 3990 bp upstream fragment of the *WER*, a 2534 bp upstream fragment of *JKD*, a 2414 bp upstream fragment of *SCR* and a 2525 bp upstream fragment of *SHR*, were introduced before HDA19-EGFP construct, respectively, as reported ^19, 49^. All the resultant constructs were verified by sequencing. Plasmid DNA was transformed into *Agrobacterium* tumefaciens strain GV3101. Plant transformation was undertaken according to A. tumefaciens-mediated floral dip transformation protocol ^50^ and the seeds harvested were screened on the MS medium with antibiotic or herbicide selection. The resistant T1 seedlings were planted in the soil for seeds collection. Either T2 seedlings or homozygous T3 or T4 seedlings were used for GFP observation, phenotype analysis and other experiments.

### Microscopy

Transverse serial sections of Arabidopsis root tips were stained with Toluidine Blue to analyze the epidermal cell differentiation ^15^. Histochemical analysis of GUS reporter lines was performed as described previously ^51^. Photographs were taken with the microscope Olymbus BX51 and SPOT-RT3 camera.

GFP and YFP expression lines were first counterstained with 20 mg/mL Propidium Iodide (PI) for 1 to 2 min and observed with the Zeiss LSM 710 NLO & DuoScan System. GFP and PI signals were observed on separate channels, with excitation at 488 nm, detection between 493 and 542 nm for GFP and excitation at 561 nm, detection between 566 and 628 nm for PI. At least 10 roots of 8-day-seedlings were examined for each line.

### RNA Isolation and qRT-PCR

Total RNA was extracted from root tips of 5 day-old or 8 day-old seedlings using the RNeasy Plant Mini kit (Qiagen) according to the manufacturer’s instructions. After treatment with DNase I (Promega), complementary DNA was synthesized from 2 mg of total RNA using the SuperScript III Reverse Transcriptase kit (Invitrogen). The qRT-PCR was performed with the Applied Biosystems 7500 Real-Time PCR System using TaKaRa SYBR Premix Ex Taq. Relative gene expression was calculated using the ΔΔCt method ^52^ using *GAPDH* as an internal control. Gene-specific primers used for RT-qPCR are listed in Supplemental Table S2.

### ChIP Assays

ChIP assays for detection of histones H3 and H4 acetylation level were carried out as described previously ^53^. Root tips were cut from 8 day-old seedlings and treated with formaldehyde. Extracted chromatin was sheared to an average length of 500 bp by sonication and immunoprecipitated with specific antibodies: anti-acetylated histone H3K9K14 (06-599; Millipore) and anti-acetylated histone H4K5K8K12K16 (06-866; Millipore). The enriched DNA was analyzed using quantitative real-time PCR.

ChIP assays for detection of HDA19 binding were carried out mainly according to ^54^ but with some modifications. Root tips of 8 day-old seedlings of Col, *HDA19pro:HDA19-EGFP /hda19* complementation line and *HDA19pro:HDA19-EGFP /hda19 /scr* line were harvested and treated with 10 mM dimethyl adipimidate (DMA) in phosphate buffer according to. Then after cleanse, the samples were fixed with 1% (v/v) formaldehyde for 20 min. Chromatin was then extracted and sonicated to produce an average of 500 bp DNA fragments. The commercial anti-GFP (ab290, ABcam) antibody was used to immunoprecipitate the chromatin. The enriched DNA was analyzed by quantitative real-time PCR. The primers used for ChIP assays are listed in Supplemental Table S2.

### Yeast Two-Hybrid Assay

Yeast two-hybrid experiments were performed according to The MATCHMAKER Two-Hybrid System 2 (BD Biosciences). The full-length as well as truncated Arabidopsis HDA19 cDNA were subcloned into pGBKT7 and in frame of BD domain. And the coding sequences of candidate genes as well as the truncated SCR domains were subcloned into pGADT7 and in frame of AD domain. The bait and prey constructs were co-transformed into the yeast strain AH109. Selection for interactions and β-galactosidase activity measurement using ONPG as substrate were carried on according to the manufacture’s protocol (clontech).

### BiFC Assay

The coding sequence of YFP was fragmented into an N-terminal domain of 155 amino acids (nYFP) and a C-terminal domain of 84 amino acids (cYFP) ^55^. HDA19 and SCR coding sequence were inserted into both pSPYNE-35S and pSPYCE-35S and in frame of nYFP and cYFP. After verification, the corresponding nYFP and cYFP plasmids were bombarded into onion epidermal cells with Model PDS-1000/He Biolistic Particle Delivery System according to the manufacture’s structure (Bio-Rad). The fluorescence was observed under the microscopy 24 hours after bombardment.

### FCS Assay

All FCS measurements were carried out in 5-day old plants. Briefly, time lapse images of 256 pixels x 256 pixels were acquired as previously described in Clark et al 2016 on a Zeiss 780 using a 40 X 1.2 W objective. Data was analyzed in SimFCS software as previously described in ^24^ (Table S3).

### Accession Numbers

Sequence data from this article can be found in the Arabidopsis Genome Initiative or GenBank/EMBL data libraries under the following accession numbers: BR6OX2 (At3g30180), CPC (At2g46410), ETC1 (At1g01380), GEM (At2g22475), GL2 (At1g79840), HDA6 (At5g63110), and HDA18 (At5g61070), HDA19 (At4g38130), JKD (At5g03150), MGP (At1g03840), RLK (At5g67280), SCM (At1g11130), SCR (At3g54220), SHR (At4g37650), TRY (At5g53200), TTG (At5g24520), WER (At5g14750).

## Acknowledgements

We would like to thank Prof. John Schiefelbein of Michigan University for gifts of marker lines. Dr. Feng Zhao for his help in discussion of experimental issues. Prof. Li-Jia Qu for his kindness in providing access to his instruments. Dr. Dong-Hui Wang for her help in daily lab management. We would like to also thank Natalie Clark for her assistance with the SimFCS software. This research was funded in part by a grant to PNB from the NIH (R01-GM043778), and by the Howard Hughes Medical Institute. This project was supported by the NSFC grants to SB (31370220) and WC (31370220).

## References

1. Dolan, L. et al. Cellular organisation of the Arabidopsis thaliana root. Development (Cambridge, England) 119, 71–84 (1993).

2. van den Berg, C., Willemsen, V., Hendriks, G., Weisbeek, P. & Scheres, B. Short-range control of cell differentiation in the Arabidopsis root meristem. Nature 390, 287–289 (1997).

3. Dolan, L. et al. Clonal relationships and cell patterning in the root epidermis of Arabidopsis. Development (Cambridge, England) 120, 2465–2474 (1994).

4. Berger, F., Hung, C.Y., Dolan, L. & Schiefelbein, J. Control of cell division in the root epidermis of Arabidopsis thaliana. Developmental biology 194, 235–245 (1998).

5. Schiefelbein, J., Kwak, S.H., Wieckowski, Y., Barron, C. & Bruex, A. The gene regulatory network for root epidermal cell-type pattern formation in Arabidopsis. J Exp Bot (2009).

6. Larkin, J.C., Brown, M.L. & Schiefelbein, J. How do cells know what they want to be when they grow up? Lessons from epidermal patterning in Arabidopsis. Annu Rev Plant Biol 54, 403–430 (2003).

7. Rerie, W.G., Feldmann, K.A. & Marks, M.D. The *GLABRA2* gene encodes a homeo domain protein required for normal trichome development in Arabidopsis. Genes & development 8, 1388–1399 (1994).

8. Galway, M.E. et al. The *TTG* gene is required to specify epidermal cell fate and cell patterning in the Arabidopsis root. Developmental biology 166, 740–754 (1994).

9. Lee, M.M. & Schiefelbein, J. WEREWOLF, a MYB-related protein in Arabidopsis, is a position-dependent regulator of epidermal cell patterning. Cell 99, 473–483 (1999).

10. Bernhardt, C. et al. The bHLH genes *GLABRA3* (*GL3*) and *ENHANCER OF GLABRA3* (*EGL3*) specify epidermal cell fate in the Arabidopsis root. Development (Cambridge, England) 130, 6431–6439 (2003).

11. Kirik, V., Simon, M., Huelskamp, M. & Schiefelbein, J. The *ENHANCER OF TRY AND CPC1* gene acts redundantly with *TRIPTYCHON* and *CAPRICE* in trichome and root hair cell patterning in Arabidopsis. Developmental biology 268, 506–513 (2004).

12. Caro, E., Castellano, M.M. & Gutierrez, C. A chromatin link that couples cell division to root epidermis patterning in Arabidopsis. Nature 447, 213–217 (2007).

13. Kwak, S.H., Shen, R. & Schiefelbein, J. Positional signaling mediated by a receptor-like kinase in Arabidopsis. Science 307, 1111–1113 (2005).

14. Kuppusamy, K.T., Chen, A.Y. & Nemhauser, J.L. Steroids are required for epidermal cell fate establishment in Arabidopsis roots. Proceedings of the National Academy of Sciences of the United States of America 106, 8073–8076 (2009).

15. Xu, C.R. et al. Histone acetylation affects expression of cellular patterning genes in the Arabidopsis root epidermis. Proceedings of the National Academy of Sciences of the United States of America 102, 14469–14474 (2005).

16. Li, D.X., Chen, W.Q., Xu, Z.H. & Bai, S.N. HISTONE DEACETYLASE6-defective mutants show increased expression and acetylation of *ENHANCER OF TRIPTYCHON AND CAPRICE1* and *GLABRA2* with small but significant effects on root epidermis cellular pattern. Plant physiology 168, 1448–1458 (2015).

17. Zhu, Y. et al. The Histone Chaperone NRP1 interacts with WEREWOLF to activate *GLABRA2* in Arabidopsis root hair development. Plant Cell 29, 260–276 (2017).

18. Liu, C. et al. HDA18 affects cell fate in Arabidopsis root epidermis via histone acetylation at four kinase genes. Plant Cell (2013).

19. Hassan, H., Scheres, B. & Blilou, I. *JACKDAW* controls epidermal patterning in the Arabidopsis root meristem through a non-cell-autonomous mechanism. Development (Cambridge, England) 137, 1523–1529 (2010).

20. Di Laurenzio, L. et al. The *SCARECROW* gene regulates an asymmetric cell division that is essential for generating the radial organization of the Arabidopsis root. Cell 86, 423–433 (1996).

21. Helariutta, Y. et al. The *SHORT-ROOT* gene controls radial patterning of the Arabidopsis root through radial signaling. Cell 101, 555–567 (2000).

22. Cui, H. et al. An evolutionarily conserved mechanism delimiting SHR movement defines a single layer of endodermis in plants. Science 316, 421–425 (2007).

23. Sabatini, S., Heidstra, R., Wildwater, M. & Scheres, B. *SCARECROW* is involved in positioning the stem cell niche in the Arabidopsis root meristem. Genes & development 17, 354–358 (2003).

24. Clark, N.M. et al. Tracking transcription factor mobility and interaction in Arabidopsis roots with fluorescence correlation spectroscopy. Elife 5 (2016).

25. Long, Y. et al. Arabidopsis BIRD Zinc Finger proteins jointly stabilize tissue boundaries by confining the cell fate regulator SHORT-ROOT and contributing to fate specification. Plant Cell 27, 1185–1199 (2015).

26. Moreno-Risueno, M.A. et al. Transcriptional control of tissue formation throughout root development. Science 350, 426–430 (2015).

27. Long, Y. et al. In vivo FRET-FLIM reveals cell-type-specific protein interactions in Arabidopsis roots. Nature 548, 97–102 (2017).

28. Baum, S.F., Dubrovsky, J.G. & Rost, T.L. Apical organization and maturation of the cortex and vascular cylinder inArabidopsis thaliana (Brassicaceae) roots. Am J Bot 89, 908–920 (2002).

29. Paquette, A.J. & Benfey, P.N. Maturation of the ground tissue of the root is regulated by gibberellin and SCARECROW and requires SHORT-ROOT. Plant physiology 138, 636–640 (2005).

30. Cui, H. & Benfey, P.N. Interplay between SCARECROW, GA and LIKE HETEROCHROMATIN PROTEIN 1 in ground tissue patterning in the Arabidopsis root. Plant J 58, 1016–1027 (2009).

31. Cui, H. & Benfey, P.N. Cortex proliferation: simple phenotype, complex regulatory mechanisms. Plant Signal Behav 4, 551–553 (2009).

32. Chen, W.Q., Li, D.X., Zhao, F., Xu, Z.H. & Bai, S.N. One additional histone deacetylase and 2 histone acetyltransferases are involved in cellular patterning of Arabidopsis root epidermis. Plant Signal Behav 11, e1131373 (2016).

33. Brady, S.M. et al. A high-resolution root spatiotemporal map reveals dominant expression patterns. Science 318, 801–806 (2007).

34. Alinsug, M.V., Yu, C.W. & Wu, K. Phylogenetic analysis, subcellular localization, and expression patterns of RPD3/HDA1 family histone deacetylases in plants. BMC Plant Biol 9, 37 (2009).

35. Petrasek, Z. & Schwille, P. Precise measurement of diffusion coefficients using scanning fluorescence correlation spectroscopy. Biophys J 94, 1437–1448 (2008).

36. Digman, M.A. & Gratton, E. Lessons in fluctuation correlation spectroscopy. Annual review of physical chemistry 62, 645–668 (2011).

37. Tian, L. et al. Reversible histone acetylation and deacetylation mediate genome-wide, promoter-dependent and locus-specific changes in gene expression during plant development. Genetics 169, 337–345 (2005).

38. Fong, P.M., Tian, L. & Chen, Z.J. Arabidopsis thaliana histone deacetylase 1 (AtHD1) is localized in euchromatic regions and demonstrates histone deacetylase activity in vitro. Cell Res 16, 479–488 (2006).

39. Schiefelbein, J., Huang, L. & Zheng, X. Regulation of epidermal cell fate in Arabidopsis roots: the importance of multiple feedback loops. Front Plant Sci 5, 47 (2014).

40. Dolan, L. Positional information and mobile transcriptional regulators determine cell pattern in the Arabidopsis root epidermis. J Exp Bot 57, 51–54 (2006).

41. Zhang, Y., Iakovidis, M. & Costa, S. Control of patterns of symmetric cell division in the epidermal and cortical tissues of the Arabidopsis root. Development (Cambridge, England) 143, 978–982 (2016).

42. Liu, X. et al. Transcriptional repression by histone deacetylases in plants. Mol Plant 7, 764–772 (2014).

43. Tian, L. et al. Genetic control of developmental changes induced by disruption of Arabidopsis histone deacetylase 1 (AtHD1) expression. Genetics 165, 399–409 (2003).

44. Chen, C.Y., Wu, K. & Schmidt, W. The histone deacetylase HDA19 controls root cell elongation and modulates a subset of phosphate starvation responses in Arabidopsis. Scientific reports 5, 15708 (2015).

45. Long, J.A., Ohno, C., Smith, Z.R. & Meyerowitz, E.M. TOPLESS regulates apical embryonic fate in Arabidopsis. Science 312, 1520–1523 (2006).

46. Fukaki, H. et al. Genetic evidence that the endodermis is essential for shoot gravitropism in Arabidopsis thaliana. Plant J 14, 425–430 (1998).

47. Lin, Y. & Schiefelbein, J. Embryonic control of epidermal cell patterning in the root and hypocotyl of Arabidopsis. Development (Cambridge, England) 128, 3697–3705 (2001).

48. Wada, T. et al. Role of a positive regulator of root hair development, *CAPRICE*, in Arabidopsis root epidermal cell differentiation. Development (Cambridge, England) 129, 5409–5419 (2002).

49. Kwak, S.H. & Schiefelbein, J. A feedback mechanism controlling SCRAMBLED receptor accumulation and cell-type pattern in Arabidopsis. Curr Biol 18, 1949–1954 (2008).

50. Clough, S.J. & Bent, A.F. Floral dip: a simplified method for Agrobacterium-mediated transformation of Arabidopsis thaliana. Plant J 16, 735–743 (1998).

51. Masucci, J.D. & Schiefelbein, J.W. Hormones act downstream of TTG and GL2 to promote root hair outgrowth during epidermis development in the Arabidopsis root. Plant Cell 8, 1505–1517 (1996).

52. Livak, K.J. & Schmittgen, T.D. Analysis of relative gene expression data using real-time quantitative PCR and the 2(-Delta Delta C(T)) Method. Methods 25, 402–408 (2001).

53. Saleh, A., Alvarez-Venegas, R. & Avramova, Z. An efficient chromatin immunoprecipitation (ChIP) protocol for studying histone modifications in Arabidopsis plants. Nat Protoc 3, 1018–1025 (2008).

54. Kurdistani, S.K. & Grunstein, M. In vivo protein-protein and protein-DNA crosslinking for genomewide binding microarray. Methods 31, 90–95 (2003).

55. Weinthal, D. & Tzfira, T. Imaging protein-protein interactions in plant cells by bimolecular fluorescence complementation assay. Trends in plant science 14, 59–63 (2009).

